# The generation and validation of a *de novo*-designed titin binder for use in fluorescence microscopy

**DOI:** 10.64898/2026.02.13.705800

**Authors:** Martin Rees, Andrew Beavil, Melissa Amerudin, Ay Lin Kho, Mark Pfuhl, Alicia Cuber Cabellero, Pauline Bennett, Yaniv Hinits, Heinz Jungbluth, Mathias Gautel

## Abstract

Advances in the generation of proteins *in silico* has enabled the efficient design of such that can bind to a specified target. Here, we demonstrate the use of a fluorescently-labelled *de novo*-designed protein to bind its target *in situ* and be imaged using fluorescence microscopy, a widely used experimental technique that typically relies on antibodies or similar evolutionary derived binders to identify the presence and location of targets in their native environment. Our *de novo*-designed protein binds the C-terminal domain M10 (Ig-169) of the giant muscle protein titin, which spans half a sarcomere, the basic contractile unit of striated muscle. M10 antibodies suitable for fluorescence microscopy are unavailable. Confocal microscopy of muscle sections shows the binder localises to the M-band of the sarcomere – where M10 is found – and fails to label muscle in competition experiments and in mutant muscle where M10 is absent. These results demonstrate the utility of *de novo*-designed proteins in immunostaining-like experiments and suggest a future where targets can be routinely identified in complex biological samples by *in silico*-generated binders. Such an approach avoids the need to generate antibodies or similar binders either *in vivo* or *in vitro*, which can have technical, financial and ethical challenges.

## Introduction

The recent application of machine learning to protein structure prediction and design has led to significant improvements in the ability to generate proteins *in silico* to bind to designated targets, with their use demonstrated in a range of diverse areas (see (1) for review). Although *de novo*-designed proteins have been developed as fluorophore binders for use in multiplex imaging (2) and a GFP-tagged binder has been visualised attaching to its cell-surface receptor *in vitro* (3), one area yet to be explored (to our knowledge) is the use of *de novo*-designed binders to provide information on a target’s presence and location in its native environment, analogous to how *in vivo*- and *in vitro*- generated binders are used in immunostaining experiments. While the proteins commonly used in immunostaining are antibodies or derivatives such as nanobodies (which are generated and harvested from animals, either by purifying the antibody directly from animal sera or by isolating the clonal cells and producing *in vitro*) other binders including affimers or designed ankyrin repeat proteins (DARPins) (produced using phage or ribosome display) may also be employed (4). Binders are visualised in the sample by either their direct fluorescent labelling or by addition of fluorescently labelled secondary antibodies/nanobodies that recognise the “primary” protein.

Here, we show that *de novo*-designed proteins can be employed in an analogous way to antibodies and other binders in fluorescence microscopy experiments, by binding specifically to their molecular target *in situ*. We target the C-terminal domain, M10, of the massive muscle protein titin (5), which is integral to the sarcomere, the basic contractile unit of striated muscle. With a molecular weight of approximately 3.5 MDa, a single titin molecule spans the ∼1.5 µm of a half-sarcomere, from its edge (the Z-disk) to its centre (the M-band) (6). M10 is the 169^th^ immunoglobulin domain in titin (7) and localises to the M-band (8). The repeating architecture of the sarcomere and precise disposition of titin along it makes titin an ideal protein for assessing the target specificity of *de novo*-designed proteins. There are no available antibodies raised against M10 that work in immunostaining, demonstrating that such gaps in labelling reagents can be closed by *de novo* binders.

### Brief Methods

Three *de novo* binders generated in Bindcraft (9) were recombinantly produced and their affinity for titin domain M10 was measured using isothermal titration calorimetry. The binding location of B86 on M10 was assessed by nuclear magnetic resonance. Alexa-647-labelled (FL-)B86 was added to muscle sections from mouse and human, with its localisation visualised using confocal microscopy.

## Results

Three *de novo*-designed binders to titin domain M10 generated in Bindcraft (9) were selected for *in vitro* investigation based on (i) having an interface predicted template modelling score above 0.9 and a predicted aligned error below 0.9 from Alphafold (10) predictions of the binder-target complex, and (ii) for their structural and binding site diversity. The binders, which we named Binder-86 (B86, numbered according to sequence length), B99 and B118, were expressed, purified and their binding to M10 was assessed using isothermal titration calorimetry. B86 binds M10 with a dissociation constant of 1.80 ± 0.21 µM (figure 1C, extended figure S1), similar to the domain’s natural ligands, obscurin and obscurin-like 1 (11), while B99 and B118 showed no evidence of binding (extended figure S1). 2D nuclear magnetic resonance of ^15^N-labelled M10 with B86 identified titin residues that had their spectral peaks shifted upon binding, which matched the residues predicted to be at the B86-M10 complex interface (Figure 1B, D, E).

**Figure 1.**
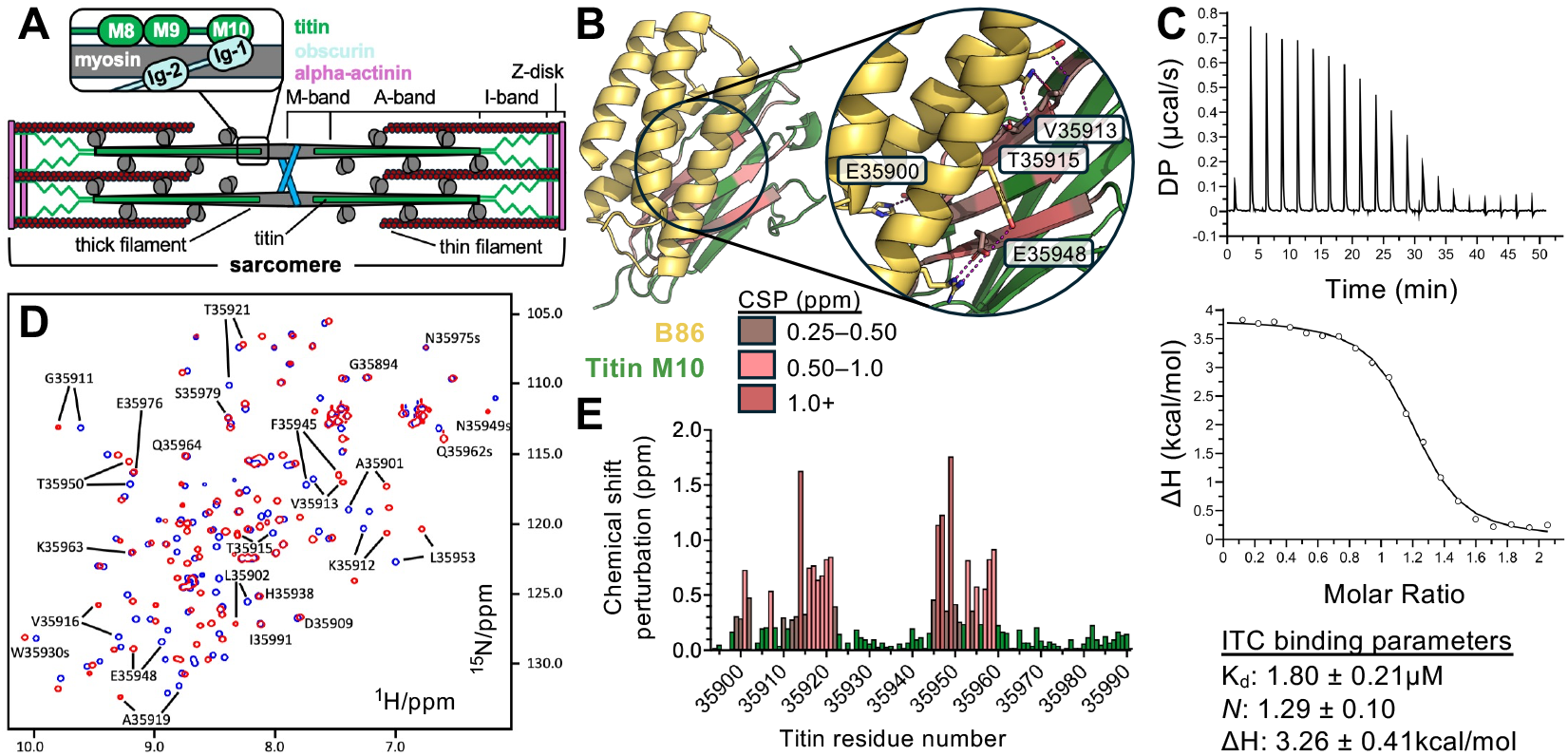
*de novo* binder B86 binds to titin domain M10. **A:** diagram of the striated muscle sarcomere. Key proteins and filaments are indicated. **B:** predicted structure of M10 (green) in complex with B86 (yellow). Chemical shift perturbations (CSP) from nuclear magnetic resonance experiments are mapped onto the M10 structure. Inset: predicted M10-B86 interface; binding residues are shown in stick format, with polar contacts indicated by magenta lines. Titin residues predicted to be make polar contacts with B86 are labelled. **C:** Representative isothermal titration calorimetry thermogram showing binding of B86 to M10 (top), with integrated data and binding model (middle) and binding parameters (bottom). **D:** Superimposed 2D ^1^H/^15^N HSQC spectra of M10 in presence (red) or absence (blue) of B86. Backbone and sidechain (“s”) cross-peaks of selected residues are labelled. **E:** Combined ^1^H and ^15^N CSP values for spectra shown in 1D. Also see extended figure S1.

M10 is found at the M-band of striated muscle sarcomeres (8). Alexa-647-Fluorescently-Labelled (FL-)B86 was added to sections isolated from mouse heart and co-incubated with an antibody raised against alpha-actinin, a protein of the sarcomeric Z-disk (12). Anti-alpha actinin was recognised by a Cy2-labelled secondary antibody. Confocal microscopy showed an alternating striated pattern (figure 2A, extended figure S2A), corresponding to localisation of anti-alpha actinin and FL-B86 at the Z-disk and M-band, respectively. To assess the specificity of the M-band labelling, FL-B86 was pre-incubated with recombinant M10 and then compared with anti-alpha actinin staining in mouse cardiac sections as before. While Z-disk staining was still present, M-band staining was reduced to a non-specific background signal (figure 2B, extended figure S2B), presumably due to sequestering of FL-B86 by excess recombinant M10.

**Figure 2.**
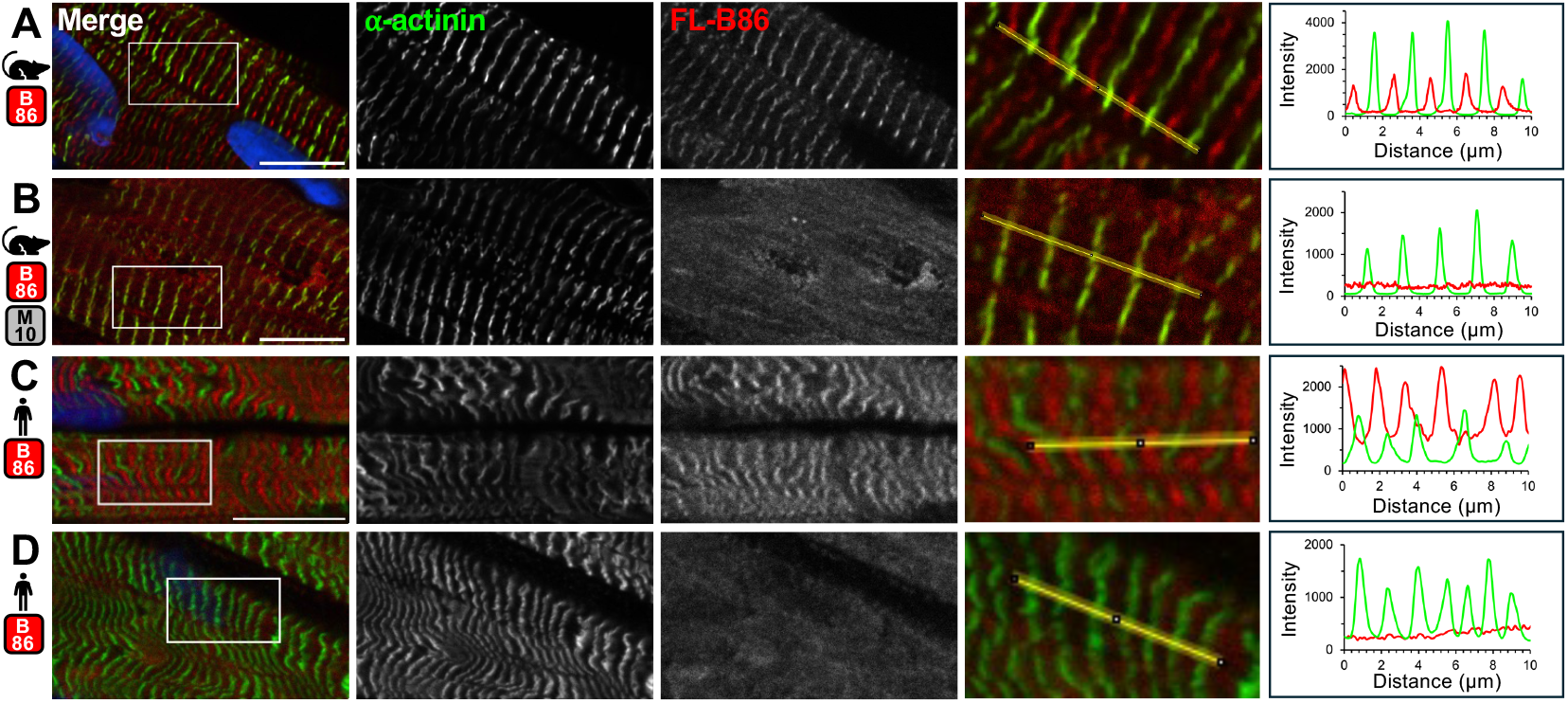
Fluorescently-labelled *de novo* binder FL-B86 localises to the striated muscle sarcomere M-band in fluorescence microscopy. **A and B:** confocal microscopy of mouse cardiac muscle sections labelled with FL-B86 (red) and anti-alpha actinin (green) in the absence (**A**) or presence (**B**) of recombinant M10. DAPI is in blue. **C and D:** confocal microscopy of human cardiac or skeletal muscle from individuals expressing (**A**) or lacking (**B**) the C-terminus of titin (including M10) labelled with FL-B86, anti-alpha actinin and DAPI. The inset shows the region selected for a line scan (yellow line) from the merged image (indicated by the white box). Colours used for the line scan are as for the images. Scale bar = 10µM. Also see extended figures S2 and S3.

We then tested the binding of FL-B86 to skeletal or cardiac muscle samples from two individuals with titin-related congenital myopathies: one carried a single titin truncating variant, whilst in the second both alleles carried titin truncating variants, meaning the former expressed full-length titin but the latter did not, lacking the final 59 kDa of the protein (patients 26 and 29, respectively, in (13)). While FL-B86 M-band staining was present in the individual expressing full-length titin (figure 2C, extended figure 3A), staining was reduced to non-specific background signal in the muscle sample lacking the C-terminus of titin (figure 2D, extended figure 3B). From this evidence, we conclude that *de-novo* binder B86 binds specifically to M10 *in situ*.

## Discussion

Here we show for the first time, to the best of our knowledge, the use of a *de novo*-designed protein to bind to its target *in situ* in fluorescence microscopy experiments, a class of ligands we call <u>F</u>luorescently <u>L</u>abelled <u>B</u>inders (FL-Bs). B86 binds to titin domain M10 *in vitro* with low-micromolar affinity and when added to muscle sections locates to the M-band of the sarcomere, where M10 is found. The specificity of the binder was demonstrated by abrogating staining when pre-incubating FL-B86 with an excess of its recombinant target, and the lack of M-band striations when staining muscle from an individual carrying two C-terminal titin truncating variants that results in the absence of M10.

With the lack of negative selection to reduce non-specific, off-target binding in their generation, it was unclear if *de novo*-designed proteins could be utilised in immunostaining-like experiments. The results presented here demonstrate they can, with FL-B86 exhibiting exquisite specificity to its target domain in the complex, protein-dense environment of striated muscle which contains at least 200 immunoglobulin domains homologous to M10 (6).

The affinity of B86 for its target is weaker than is typical for antibodies or nanobodies, but it still localises to the M-band correctly after fluorophore labelling and does not get outcompeted by endogenous ligands. The quick binding on- and off-rates of this binder could be utilised to quickly wash out the binder when performing multiplexing experiments such as cyclic immunofluorescence (14), or even be used in a way analogous to DNA-PAINT (15) with transient binding providing the fluorophore “blinking” necessary for many super-resolution microscopy techniques.

With only three *de novo* proteins tested *in vitro*, the high “hit rate” of 33% achieved here promises an affordable avenue to generate further *in situ* binders to possibly limitless molecular targets to observe using fluorescence microscopy. Their generally small size (16) aids sample penetration, access to the binders’ antigen and can increase image resolution by decreasing the linkage error (17), while the lack of *in vitro* or *in vivo* steps to binder development reduces costs, technical challenges and ethical concerns associated with animal use in research. If the promise of the *de novo* binder shown here can be repeated for other targets, we anticipate that generating fluorescently labelled binders *in silico* for immunostaining-like experiments will soon match or even surpass the generation of binders *in vivo* or *in vitro*.

## Extended Methods

### Binder generation

Binders were generated *in silico* using Bindcraft (9) version 1.5.1 installed on Ubuntu 22.04.5 LTS running on a Nvidia GeForce RTX 4080 GPU. The coordinates for titin domain M10 in its crystal structure in complex with the first immunoglobulin domain of obscurin, PDB 4C4K (18) was used as the input target structure for binder generation, with binders specified to be of length 65 – 125 amino acids, and to target either anywhere on M10 (in the case of B86) or residue H35946 (H54 in the PDB file, for B99). For B118, the input PDB was modified by mutating residues I3, E8, S15, D17, V21, A25, A27, T29, E31, E56, T58, D60, T63, I65 and M67 to lysine, and targeting residues L76, T78, T91 and N93. 100 binder sequences were produced by Bindcraft that passed its internal quality control checks for each input PDB-target combination, with a subset of the binder-target structures re-predicted using the Alphafold server (10). Models with pAE values below 0.9 and ipTM values above 0.9 were considered for *in vitro* testing, with three selected – which we named Binder-86 (B86), B99 and B118 – based upon structural and binding site diversity. Models were visualised in PyMol (Schrödinger Inc.). The selected binder sequences were:

B86

SMTPSEEILKEMIKLVEELIETEDESIKDKIMVLMYEWVRALREEGIKLPHEKVVLFAQIVRDLQMTPITDKESLKKIIEKLKSLL

B99 SEVEKFADELVKKLEEELPKRNTGNPMNWTHNVIEFFHEEVFRKRKLGWGGYIIDDGDTLHVHLIITRDTPDGSKEVLAELNIEIEVTETEAKVKSYSL

B118 SAELEARMAETIAEFLELLKELKKKWTEFLTELGYEEEKERYKKYYDHLLSLRTVEEWLAFIPRHDEFQREVYPNGHWWLWPRLNEEQQKKYVEAHEIQTKADAHLTRLDYL ARRLQE

### Protein production

pET-IDT vectors containing a DNA sequence encoding the three selected binders with an N-terminal His_6_ tag and TEV protease cleavage site (sequence MHHHHHHSTENLYFQGSS) were purchased from Integrated DNA Technologies. A pET vector encoding titin domain M10 (residues 35893 – 35991 of the Inferred Complete sequence, NM_001267550 (19)) with the same N-terminal tag sequence was generated by amplifying the sequence from a cardiac cDNA library and recombined into the vector by Fast Cloning (20). All vectors were transformed into (DE3)-BL21 RIPL cells (Agilent) and cultured in either Luria Bertani media (for binders and M10) or minimal media containing ^15^N-labelled glucose (for M10 used in NMR) with antibiotics, and protein expression was induced by addition of IPTG to 0.5 mM, followed by overnight incubation at 18°C with shaking at 200 RPM. Cultures were harvested by centrifugation, resuspended in buffer A (30 mM HEPES pH 7.0, 300 mM NaCl, 20 mM imidazole) with a cOmplete protease inhibitor tablet (Sigma) and lysed by addition of lysozyme and B-Per (ThermoFisher), with DNA sheared by sonication. The soluble fraction was separated from the insoluble fraction by centrifugation at 17,000g for 40 minutes and the recombinant protein was purified by nickel affinity chromatography using a 1 mL HisTrap column (GE Healthcare), eluted with buffer B (buffer A with 500 mM imidazole) followed by size-exclusion chromatography (SEC) on a Superdex-75 (GE Healthcare) equilibrated in 20 mM HEPES pH 7.0, 100 mM NaCl. For ^15^N-labelled M10, the His_6_ tag was cleaved using HyperTev60 (21), the sample passed through a second HisTrap column and then loaded onto a Superdex-75. Elution fractions from SEC containing pure recombinant protein were collated and concentrated for downstream use.

### Isothermal titration calorimetry

*De novo* binders at 0.2 – 1 mM were injected from the syringe into the cell containing 20 – 100 µM titin M10 or buffer only (20 mM HEPES pH 7.0, 100 mM NaCl) at 25°C, with 19 injections of 2 µl in total on a MicroCal PEAK-ITC (Malvern). Data points of the heat evolved were corrected with a fitted offset, and binding and thermodynamic parameters were calculated by fitting a single-site binding model in PEAQ-ITC Analysis (Malvern). For B86 with M10, *n* = 3 technical repeats.

### Nuclear magnetic resonance

B86 and ^15^N M10 were dialysed into 20 mM sodium phosphate pH 7.0, 50 mM NaCl, 2 mM DTT, 0.02% sodium azide. 1H-15N HSQC experiments (hsqcfpf3gpphwg) provided by the manufacturer (Bruker) with a combination of 1H flip back pulses centred on the water resonance plus watergate (22) were recorded on samples of 200 µM 15N-labelled M10 using a Bruker Neo 800 MHz spectrometer (equipped with a TCI cryoprobe) in the presence or absence of 400 µM unlabelled B86. Spectra were processed with Bruker Topspin 4.5 to a size of 2048 * 2048 points with a qsine window function using a shift of 2 in both dimensions. Spectra were analysed using CCPNMR 3.3.4 (23) including peak picking, assignment and calculation of chemical shift perturbations (CSP). Assignments were taken from BMRB entry 25305 (24).

### Protein labelling

500 µg B86 in PBS was incubated with 53 µg Alexa Fluor 647 NHS ester (ThermoFisher) for 1 hour at room temperature, with the reaction quenched by addition of Tris pH 7 to 0.1 M, then the protein-dye conjugate (Fluorescently-Labelled B86, FL-B86) was separated from the free dye by passing through a Nap-5 (G25) column (GE) pre-equilibrated in PBS. Fractions containing the protein-dye conjugate (at a protein:dye ratio of 1:0.4) were collated and concentrated to ∼1 mg/mL.

### Immunostaining

10 µm sections of snap-frozen tissue (mouse hearts and human biopsies) were prepared using a CM1950 cryostat (Leica Biosystems), then permeabilised for 2 minutes with 0.2% saponin and blocked with 5% normal goat serum in 1% BSA dissolved in Gold Buffer (1 mM Tris-HCl pH 7.5, 155 mM NaCl, 2 mM MgCl, 2 mM EDTA) for 1 hour at RT. Sections were incubated with anti-alpha actinin (Sigma-Aldrich A7811, 1:500 dilution), FL-B86 (1.46 µM) and optionally M10 (14.6 µM) diluted in 1% BSA/Gold Buffer overnight at 4°C, incubated with DAPI and Cy2 AffiniPure goat anti-mouse IgG (Jackson ImmunoResearch 115-225-146, 1:100 dilution) diluted in 1% BSA/Gold Buffer for 1 hour at RT and then mounted with mounting medium (30 mM Tris-HCl pH 9.5, 70% v/v glycerol and 236 mM *n*-propyl gallate).

### Microscopy

Confocal microscopy images were acquired on a Nikon AXR inverted confocal microscope equipped with a 60 X or 100 X objective at the KCL Nikon Imaging Centre and prepared in FIJI (25), with line scans of pixel intensities of the anti-alpha actinin and FL-B86 channels measured using the “Plot Profile” function.

### Ethics

Animal husbandry and experimental procedures were carried out according to UK Home Office guidelines (Animals Scientific Procedures Act 1986) under project licence PP3311131. Patients 26 and 29 from (13) were enrolled under appropriate procedures in accordance with the ethical guidelines from their local institutions, with written consent obtained from them or their legal guardians for genetic studies. The study was conducted in accordance with approvals by the National Research Ethics Service (NRES) London – West London & GTAC (Reference: 06:Q0406/33).

## Funding and acknowledgements

M.R, M.A, A.L.K and M.G are supported by the British Heart Foundation (grants RG/F/22/110079, FS/4yPhD/F/23/34199 and CH/08/001). M.R was supported by the Randall Centre pump-prime funding scheme. A.C.C is supported by the King’s BIPAS CDT award (ST12909). Y.H is supported by the European Research Council (856118). NMR and ITC data were collected at the KCL Centre for Biomolecular Spectroscopy; we thank Agrim Gupta and Tam Bui for their help. We thank the Nikon Imaging Centre at King’s College London for help with confocal microscopy.

## Author Contributions

M.R: Conceptualization (lead), funding acquisition, investigation, project administration, supervision, visualization, writing – original draft preparation. A.B: conceptualization, investigation, writing – review and editing. M.A, A.L.K, M.P, A.C.C, P.B, Y.H: investigation, writing – review and editing. H.J: resources, writing – review and editing. MG: funding acquisition, writing – review and editing.

---

**Extended Data Figure S1.**
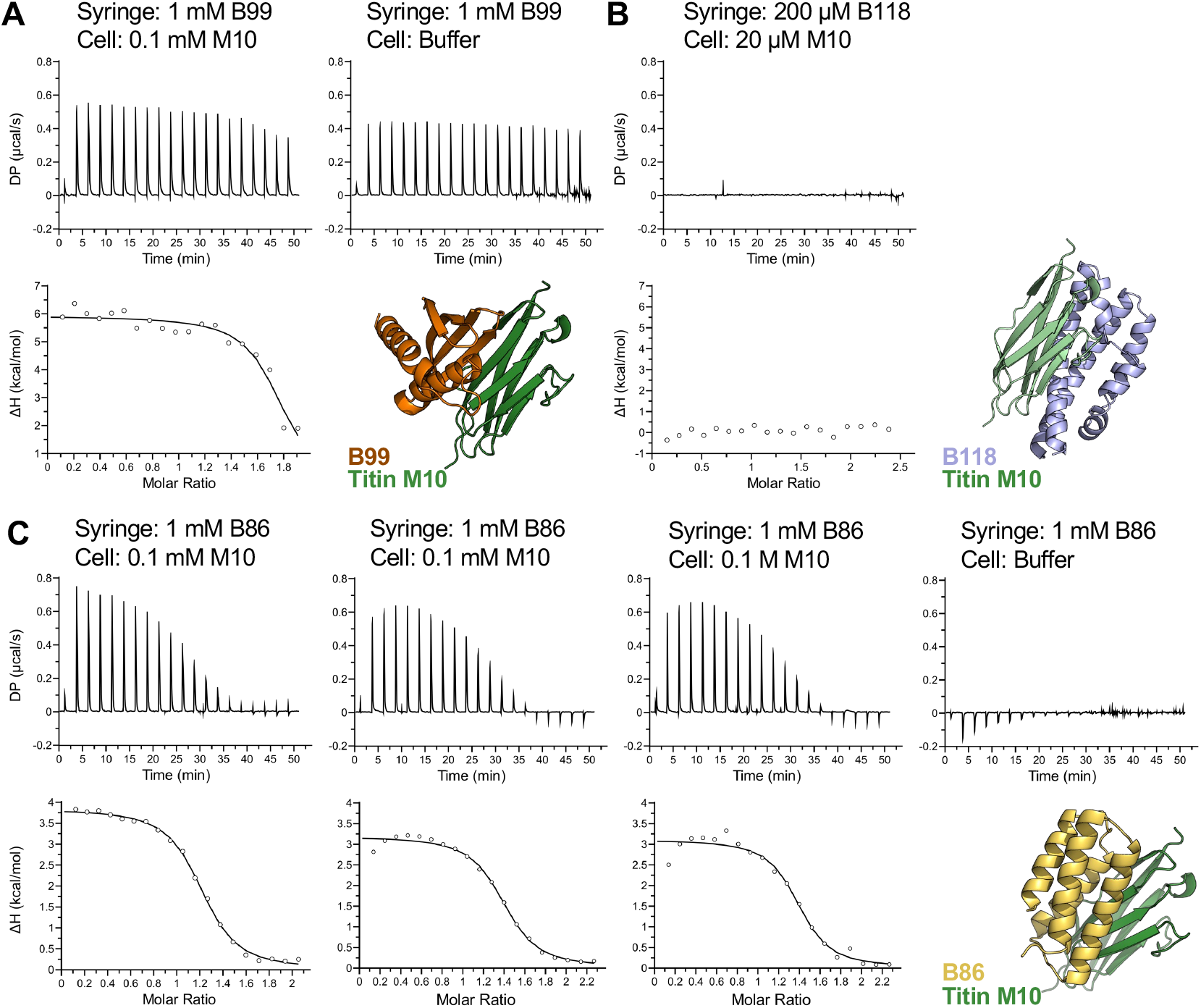
Isothermal titration calorimetry experiments testing the binding of *de novo* binders B86, B99 and B118 to titin M10. **A:** B99. **B:** B118. **C:** B86. For each experiment, the thermogram is showed at (top) and the integrated data at (bottom) with single-site binding model fitted if possible. Injection of B99 and B86 into buffer-only shows the heat evolved upon dilution of the binder. The predicted binder:M10 structures are shown for each experiment.

**Extended Data Figure S2.**
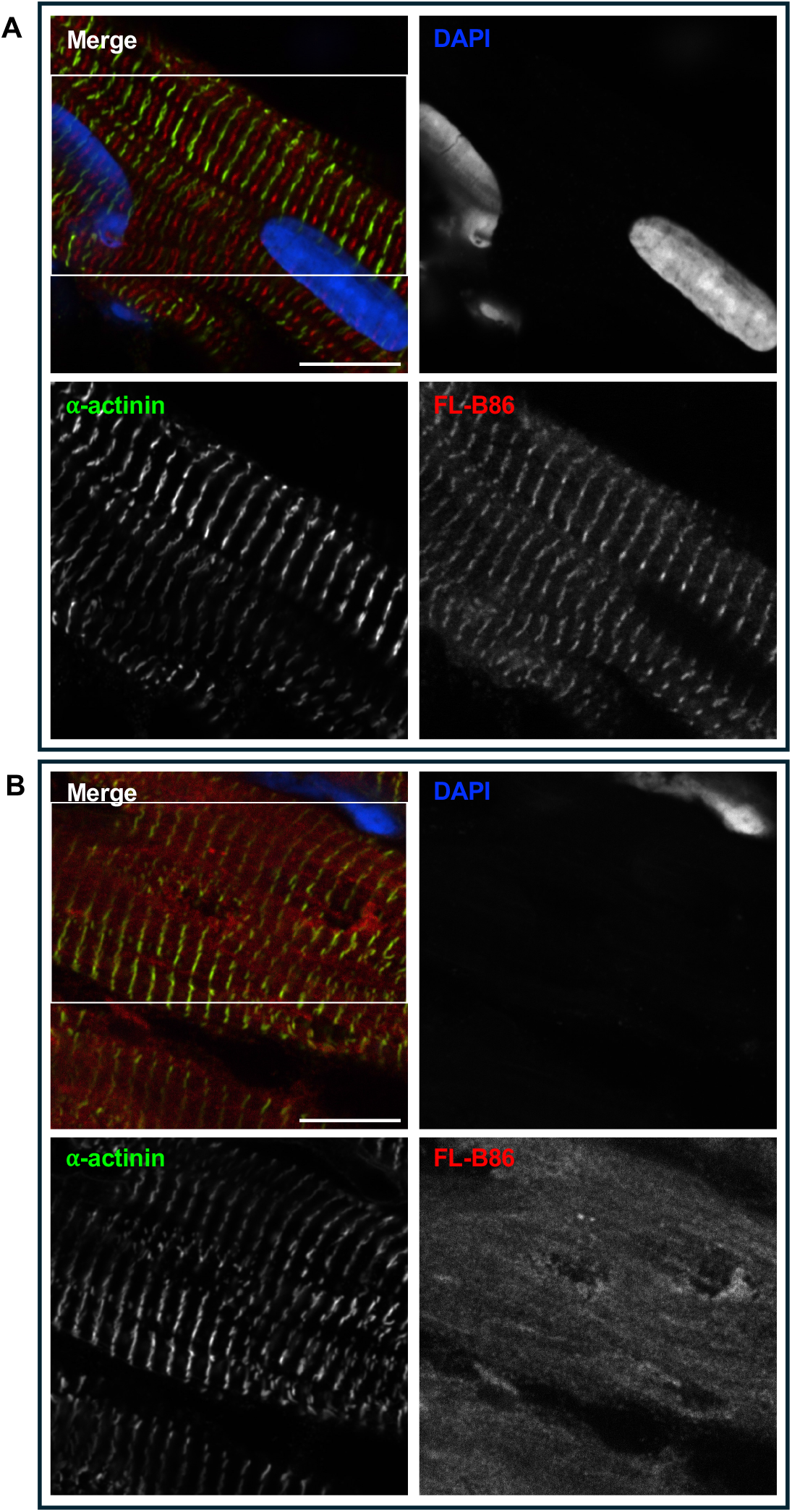
Full-size confocal images of mouse cardiac muscle labelled with anti-alpha actinin (green), FL-B86 (red) and DAPI (blue). **A:** in absence of recombinant M10 (as shown in Figure 2A). **B:** binder mixture incubated with recombinant M10 prior to muscle labelling (as shown in Figure 2B). Scale bar = 10µM. Area shown in Figure 2 indicated on merged image.

**Extended Data Figure 3.**
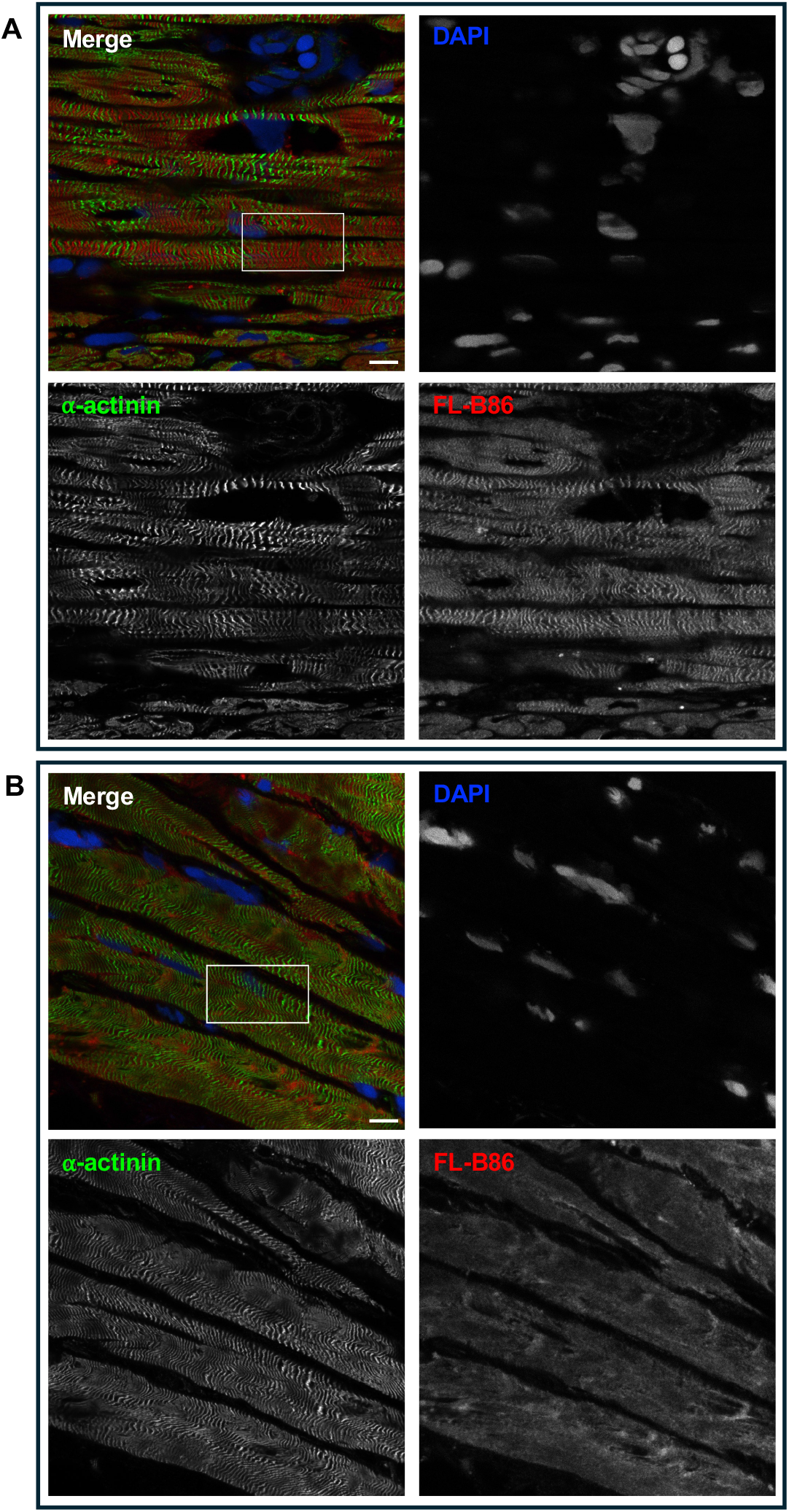
Full-size confocal images of human cardiac or skeletal muscle from individuals with titin truncating variants labelled with anti-alpha actinin (green), FL-B86 (red) and DAPI (blue). **A:** patient 26 from Rees *et al* 2021 containing 1 complete titin transcript **B:** patient 29 from Rees *et al* 2021 containing no complete titin transcripts, and lacking M10. Scale bar = 10µM. Area shown in Figure 2 indicated on merged image.

